# Fine-scale recombination landscapes between a freshwater and marine population of threespine stickleback fish

**DOI:** 10.1101/430249

**Authors:** Alice F. Shanfelter, Sophie L. Archambeault, Michael A. White

## Abstract

Meiotic recombination is a highly conserved process that has profound effects on genome evolution. Recombination rates can vary drastically at a fine-scale across genomes and often localize to small recombination “hotspots” with highly elevated rates surrounded by regions with little recombination. Hotspot targeting to specific genomic locations is variable across species. In some mammals, hotspots have divergent landscapes between closely related species which is directed by the binding of the rapidly evolving protein, PRDM9. In many species outside of mammals, hotspots are generally conserved and tend to localize to regions with open chromatin such as transcription start sites. It remains unclear if the location of recombination hotspots diverge in taxa outside of mammals. Threespine stickleback fish *(Gasterosteus aculeatus)* are an excellent model to examine the evolution of recombination over short evolutionary timescales. Using an LD-based approach, we found recombination rates varied at a fine-scale across the genome, with many regions organized into narrow hotspots. Hotspots had divergent landscapes between stickleback populations, where only ~15% were shared, though part of this divergence could be due to demographic history. Additionally, we did not detect a strong association of PRDM9 with recombination hotspots in threespine stickleback fish. Our results suggest fine-scale recombination rates may be diverging between closely related populations of threespine stickleback fish and argue for additional molecular characterization to verify the extent of the divergence.

## Introduction

Meiotic recombination is a highly-conserved process across a broad range of taxa (de Massy 2013; Petes 2001). Recombination creates new allelic combinations by breaking apart haplotypes (Coop and Przeworski 2007; Otto and Lenormand 2002), promotes the proper segregation of chromosomes during meiosis in many species (Davis and Smith 2001; Fledel-Alon et al. 2009; Kaback et al. 1992; Mather 1936), and has a pronounced impact on the evolution of genomes (Mugal et al. 2015; Webster and Hurst 2012). In many species, meiotic recombination occurs in small 1–2 kb regions called recombination “hotspots” which are surrounded by large genomic regions with little to no recombination (Barton et al. 2008; Baudat et al. 2010; Hellsten et al. 2013; Jeffreys et al. 1998; McVean et al. 2004; Myers et al. 2005; Steiner et al. 2002).

In most species, hotspot location is highly conserved over long evolutionary timescales (Kawakami et al. 2017; Lam and Keeney 2015; Singhal et al. 2015; Tsai et al. 2010). For example, finches share upwards of 73% of hotspots across 3 million years of evolution (Singhal et al. 2015) while species of *Saccharomyces* share 80% of hotspots over 15 million years of evolution (Lam and Keeney 2015). Evolutionarily conserved hotspots are often localized around regions of open chromatin such as transcription start sites (TSSs) and CG-rich regions (i.e. CpG islands) in vertebrates (Auton et al. 2013; Kawakami et al. 2017; Lee et al. 2004; Pan et al. 2011; Pokholok et al. 2005; Singhal et al. 2015; Tischfield and Keeney 2012). This localization pattern is thought to be due the opportunistic nature of Spo11, a meiosis specific protein which initiates recombination by creating double stranded breaks at regions of open chromatin (Celerin et al. 2000; Ohta et al. 1994; Pan et al. 2011).

A notable exception to strong conservation of recombination hotspots has been documented in mammals, where hotspot location evolves rapidly between closely related species or even between populations (Baker et al. 2015; Brick et al. 2012; Pratto et al. 2014; Smagulova et al. 2016; Stevison et al. 2015). Contrary to the pattern observed in conserved systems, rapidly evolving hotspots typically form away from functional genomic elements and are localized by the zinc finger histone methyltransferase protein, PRDM9 (Baker et al. 2015; Baudat et al. 2010; Billings et al. 2013; Brick et al. 2012; McVean et al. 2004; Myers et al. 2005; Myers et al. 2010; Myers et al. 2008; Parvanov et al. 2010; Powers et al. 2016; Pratto et al. 2014). PRDM9 contains multiple DNA-binding zinc fingers that are under strong positive selection, leading to divergent hotspot localization between closely related species (Baker et al. 2015; Billings et al. 2013; Brick et al. 2012; Myers et al. 2010; Parvanov et al. 2010). Though rapidly evolving hotspots have only been documented in some mammals, positive selection is acting on the zinc finger domain of PRDM9 orthologs in many non-mammalian species (Baker et al. 2017; Oliver et al. 2009). This raises the intriguing possibility that some species outside of mammals may also have rapidly evolving hotspots. It is also possible that PRDM9 is not necessary for rapid evolution of hotspots in other species and that other mechanisms could lead to the evolution of fine-scale rates of recombination over short timescales.

Threespine stickleback fish *(Gasterosteus aculeatus)* are an excellent system to study the evolution of fine-scale recombination rates. Multiple populations of threespine stickleback fish have independently adapted to freshwater environments from marine ancestors in the last 10–15 thousand years (Bell and Foster 1994; Orti et al. 1994), providing the opportunity to study the parallel evolution of hotspots in well-characterized populations across the Northern Hemisphere (Bell and Foster 1994; Ostlund-Nilsson et al. 2007; Wootton 1976). Broad-scale recombination rates have been examined in threespine stickleback using genetic crosses (Glazer et al. 2015; Peichel et al. 2001; Roesti et al. 2013; Sardell et al. 2018), but fine-scale recombination rates have not been estimated due to low marker density.

Fine-scale recombination rates can be estimated through a variety of approaches. Recombination rates can be directly measured through genetic linkage maps (Broman et al. 1998; Campbell et al. 2016; Drouaud et al. 2006; Marand et al. 2017) or though sperm genotyping (Baudat and de Massy 2007; Guillon and de Massy 2002; Jeffreys et al. 2001). Both methods require a large number of progeny or sperm and a high density of genetic markers to capture a sufficient number of crossovers. Recombination rates can also be indirectly measured by identifying the binding sites of proteins that initiate double strand breaks (Pratto et al. 2014; Smagulova et al. 2011) as well as repair double strand breaks through homologous recombination (Dumont and Payseur 2011; Froenicke et al. 2002). Another broadly used approach estimates recombination rates from patterns of linkage disequilibrium (LD) in populations, providing a historical measure of meiotic crossovers over multiple generations (Chan et al. 2012; McVean et al. 2004; Myers et al. 2005; Wall and Stevison 2016). LD-based methods are able to estimate rates at a fine-scale, but rate estimation can be biased by demographic history (e.g. bottlenecks, population expansions, population sub-structure, etc.) (Dapper and Payseur 2017; Johnston and Cutler 2012), increasing false negative and false positive rates when calling recombination hotspots (Dapper and Payseur 2017). Despite the higher error rates, many recombination hotspots identified through LD-based methods have been validated using other approaches (Jeffreys et al. 2005; Morgan et al. 2017; Myers et al. 2006).

Here, we used an LD-based approach to estimate genome-wide recombination rates in a marine (Puget Sound) and freshwater (Lake Washington) population of threespine stickleback fish. We found recombination landscapes varied at a fine-scale between the two populations, often organized into recombination hotspots. We found most recombination hotspots were not shared between populations. We describe how the complex demographic histories of threespine stickleback fish populations (Bell and Foster 1994; Ferchaud and Hansen 2016; Hohenlohe et al. 2010; Liu et al. 2016) may influence the overall distribution of recombination hotspots and argue that the patterns we observe may not be completely driven by population bottlenecks. Additionally, we found little evidence that threespine stickleback hotspots are associated with PRDM9 binding, indicating hotspots are likely localized by a different mechanism.

## Materials and Methods

### Whole genome sequencing and assembly

Genomic DNA was extracted from caudal tail clips of 13 female and 12 male fish collected from Lake Washington (freshwater population; Washington, USA) and 18 female and 6 male fish collected from Northern Puget Sound (marine population; Washington, USA) using a standard phenol-chloroform extraction. Paired-end libraries were prepared using the Illumina TruSeq kit and were size-selected to target 400 bp fragments. Libraries were multiplexed and sequenced on Illumina NextSeq lanes for 300 cycles (Georgia Genomics and Bioinformatics Core, University of Georgia). Residual adapter sequences and low quality regions were trimmed from the sequencing reads using Trimmomatic (v0.33) with the following parameters: PE –phred 33 slidingwindow:4:20. Trimmed reads were aligned to the revised threespine stickleback genome assembly (supplemental file S5, https://datadryad.org/resource/doi:10.5061/dryad.q018v/1) (Glazer et al. 2015) using Bowtie2 (v2.2.4, default parameters) (Langmead and Salzberg 2012). With these parameters, the average alignment rate for Lake Washington was 94.2% and 87.3% for Puget Sound. Reads with a mapping PHRED quality score of 20 or less were removed from the analysis (Samtools, v1.2.0, default parameters) (Li et al. 2009). For Puget Sound, four female individuals had 5x or lower sequencing coverage and were removed from the analysis. After removing poorly aligned reads and low coverage individuals, the average read coverage across all individuals in each population was 17x and 22x for Lake Washington and Puget Sound, respectively.

Two outgroup species were used to infer ancestral allele states and to estimate mutation matrices for each population (see Estimation of Recombination Rates). Whole-genome Illumina sequences for one female ninespine stickleback fish *(Pungitius pungitius*, DRX012173) (White et al. 2015) and one female blackspotted stickleback fish *(Gasterosteus wheatlandi*, DRX012174) (Yoshida et al. 2014) were aligned to the revised threespine stickleback genome assembly (Glazer et al. 2015) using Bowtie2 (v2.2.4). Less stringent alignment parameters were used to allow for greater sequence divergence between threespine stickleback and each outgroup (-D 20 −R 3 −N 1 −L 20 −I S,1,0.50 -rdg 3,2 -rfg 3,2 -mp 3). The overall alignment rate of *P. pungitius* was 46.0% whereas the overall alignment rate of *G. wheatlandi* was 74.2%. The higher alignment rate of *G. wheatlandi* is consistent with *G. wheatlandi* sharing a more recent common ancestor with *G. aculeatus* (Kawahara et al. 2009).

### SNP genotyping

Single nucleotide polymorphisms (SNPs) were genotyped in each threespine stickleback population and outgroup species independently following the GATK best practices for SNP discovery for whole genome sequences (v3.6) (Van der Auwera et al. 2013). PCR duplicates were removed using MarkDuplicates (REMOVE_DUPLICATES=true). Regions around insertions or deletions (indels) were realigned with RealignerTargetCreator (default parameters) and IndelRealigner (default parameters). Variants were called for each individual using HaplotypeCaller (genotyping mode DISCOVERY). Joint genotyping (GenotypeGVCFs, default parameters) was completed by pooling all individuals for each population. Low-quality SNPs were filtered from the data set using vcftools (v0.1.12b) (Danecek et al. 2011) with the following filters: removing all sites with more than two alleles, removing sites where genotype data was missing among individuals, removing sites where the population mean depth coverage was less than half or greater than twice the average coverage for each population (Lake Washington: 8x – 24x read depth coverage; Puget Sound: 11x - 44x read depth coverage), and removing sites with a genotype quality score less than 30. Singletons and sites fixed for the alternate allele across all individuals in a population were also removed. After filtering, the Lake Washington population had 5,054,729 SNPs genome-wide (11 SNPs/kb) and the Puget Sound population had 4,142,876 SNPs (9 SNPS/kb) genome-wide (prior to filtering Lake Washington had 11,937,220 SNPs and Puget Sound had 11,070,421 SNPs). For the outgroup species, *P. pungitius* and *G. wheatlandi*, low-quality SNPs were excluded by removing variants with a genotyping quality score less than 30 or a read depth less than two, resulting in 13,691,521 SNPs genome-wide in *G. wheatlandi* (16,783,618 SNPs prior to filtering) and 7,791,420 in *P. pungitius* (26,173,287 SNPs prior to filtering).

### Haplotype phasing

Each chromosome was phased independently with SHAPEIT (v2.r837), a read-aware phasing tool (Delaneau et al. 2013). Phase-informative reads with two heterozygous SNPs on the same read were identified to assist with the estimation of haplotypes. Phase-informative reads had a mapping quality score greater than 20. Convergence of the MCMC algorithm was estimated by examining switch error rates between individual runs. A low switch error rate would indicate that the MCMC phasing runs have converged on a similar haplotype configuration. Switch error was measured using vcftools (v0.1.12b) using –diff-switch-error (Danecek et al. 2011). A low switch error was achieved within a reasonable run time with the following SHAPEIT parameters: --main 2000 --burn 200 --prune 210 --states 1000 (average switch error between phasing runs: 0.824% for Lake Washington and 1.26% for Puget Sound). All other parameters were left at the default values.

### Estimation of recombination rates

Recombination rates were estimated with LDHelmet (v1.7) (Chan et al. 2012). LDHelmet estimates historical recombination rates from population data by analyzing patterns of linkage disequilibrium across phased individuals. The ancestral allele state was defined for every SNP in each threespine stickleback population by comparing to the allele present in the two outgroup species. An ancestral allele state could not be assigned if a polymorphism was segregating among the outgroup species. Therefore, SNPs were only assigned an ancestral state if *P. pungitius* and *G. wheatlandi* were homozygous for the same allele. The ancestral allele was assumed to be the nucleotide carried by *P. pungitius* and *G. wheatlandi*, and was assigned a prior probability of 0.91. To allow for uncertainty in the ancestral allele state, the other three possible nucleotides were assigned prior probabilities of 0.03. If the ancestral allele state could not be inferred, the prior probability of each nucleotide being the ancestral allele was set as the overall frequency of that particular nucleotide on the chromosome. Nucleotide frequencies were empirically determined from all sites on a threespine stickleback chromosome where *P. pungitius* and *G. wheatlandi* had read coverage that passed the filtering scheme. Mutation matrices were estimated for each population separately. For every position where an ancestral allele state could be inferred, the total number of each type of mutation away from the ancestral allele was quantified. A normalized 4×4 mutation matrix was generated for each chromosome as previously described (Chan et al. 2012). The ancestral allele state and mutation matrices were generated using a custom Perl script.

Each LDHelmet module was run using the following parameters. Custom Python scripts were used to create the SNP sequence and SNP position input files. Full FASTA sequence were created using vcf2fasta from vcflib (available at https://github.com/vcflib/vcflib). Haplotype configuration files were created for each chromosome with the find_confs module using a window size of 50 SNPs (-w 50). Likelihood tables were created using table_gen with the recommended grid of population scaled recombination rates per base pair (ρ/bp) (-r 0.0 0.1 10.0 1.0 100.0). Watterson’s θ was estimated using a custom Python script with the R package, PopGenome (Pfeifer et al. 2014), where Watterson’s θ was calculated in 2 kb regions with a sliding window of 1 kb and all windows were averaged together. To maintain a reasonable computational time, a single representative likelihood lookup table was generated for the autosomes of each population from chromosome one, using the average Watterson’s θ between Lake Washington and Puget Sound (-t 0.002). Although Watterson’s θ was different between the Lake Washington and Puget Sound populations, previous studies have determined that small changes to parameters such as Watterson’s θ do not affect the final likelihoods (Auton and McVean 2007; McVean et al. 2004). Separate likelihood tables were created for the pseudoautosomal region of the sex chromosomes (chromosome 19). Padé coefficient files were created using the module pade with a Watterson’s θ of 0.002 and the recommended 11 padé coefficients (-t 0.002 −x 11). The module rjmcmc was run for 1 million iterations with 100,000 burn in iterations, a block penalty of 10, and a window size of 50 SNPs (-w 50 -b 10 -burn_in 100000 -n 1000000). Population-scaled recombination rates were extracted from the rjMCMC run with the post_to_text module. Recombination rates were reported in ρ/bp where p is a population scaled recombination rate (4N_e_r).

### Correlation with genetic maps

Population-scaled recombination rates were compared with recombination rates estimated from a high-density genetic linkage map (Glazer et al. 2015). Recombination rates from LDHelmet were converted from ρ/bp to cM/Mb as previously described (Smukowski Heil et al. 2015). Briefly, the recombination rate (cM/Mb) was calculated between every pair of adjacent markers in the genetic map and a chromosome-wide recombination rate was calculated as the average among the regions. The average LD-based recombination rate (ρ/Mb) was computed in the same individual regions of a chromosome in Lake Washington and Puget Sound by averaging the per bp rho estimate across the total length of the region (ρ/Mb). A single conversion factor was calculated for each chromosome. Each conversion factor was calculated by dividing the average linkage map recombination rate for a chromosome (in cM/Mb) by the average LD-based recombination rate (ρ/Mb) for that chromosome.

### Identification of recombination hotspots

Recombination hotspots were defined using a sliding window approach. In each window, the average recombination rate within a 2 kb window was compared to the average recombination rate from the 40 kb regions flanking either side of the 2 kb window. Hotspots were defined as the 2 kb regions that had a 5-fold or higher recombination rate relative to the mean recombination rate in the flanking background regions. The 2 kb windows iterated forward in 1 kb increments. If multiple hotspots were found within a 5 kb region, only the hotspot with the highest rate was retained. Misassemblies in the reference genome could generate false hotspots. To limit this, all hotspots that spanned a contig boundary in the reference genome were removed (384 hotspots out of 4,349 total hotspots). Hotspots were considered shared between populations if the midpoints of the two hotspots were within 3 kb of each other. Random permutations were used to calculate the expected amount of hotspot overlap between Lake Washington and Puget Sound. 10,000 random permutations were drawn from the genome totaling the number of 2 kb hotspots for each population. Recombination hotspots were identified and filtered using custom Perl and Python scripts.

### Genetic variation within and between populations

Within population nucleotide diversity (π) and Tajima’s D were calculated separately for each chromosome. To capture rare variants, previously excluded singletons were included in the analysis. Nucleotide diversity and Tajima’s D were calculated using the R package, PopGenome (Pfeifer et al. 2014) and a custom Python script. Nucleotide diversity was calculated between populations by combining SNP variants among all individuals in each population. Population structure was estimated between Lake Washington and Puget Sound using FastStructure (v1.0) (Raj et al. 2014). For this analysis, SNPs from Lake Washington and Puget Sound were merged using vcftools (Danecek et al. 2011) and only biallelic sites with no missing data were retained. The sex chromosomes (chromosome 19) were also excluded. The final SNP dataset was composed of 4,113,937 SNPs. Three trials were completed at K values of 1, 2, and 3. These K values were chosen to differentiate scenarios where Lake Washington and Puget Sound were one panmictic population (K=1) or Lake Washington and Puget Sound were two distinct populations (K=2). A K of 3 was chosen to identify any hidden population structure within either population. The model that best explained the population structure was determined using chooseK.py (Raj et al. 2014) and the structure plot was visualized using distructK.py (Raj et al. 2014).

### Estimation of demographic history

Demographic history can affect LD-based estimates of recombination rates (Dapper and Payseur 2017; Johnston and Cutler 2012). To determine whether the demographic history of threespine stickleback fish could influence the ability to detect recombination hotspots, hotspots were assayed in simulated haplotypes with known recombination profiles and demographic histories. Demographic histories used in the simulations were based on the estimated histories of Lake Washington and Puget Sound, modeled using a Pairwise Sequentially Markovian Coalescent (PSMC) process with default parameters (Li and Durbin 2011; Liu and Hansen 2017). PSMC was run on one female from Lake Washington and one female from Puget Sound. Confidence intervals were estimated on 100 bootstrap replicates. Demographic histories were visualized using psmc_plot.pl (Li and Durbin 2011).

### Simulations using estimated demographic histories

Using the demographic histories estimated with PSMC, 250 kb haplotypes with four 2 kb recombination hotspots were simulated using the program fin, part of the LDHat software package (Auton and McVean 2007; McVean et al. 2004). The hotspots were place 50 kb apart at 75, 125, 175, and 225 kb. The background recombination rate was set at 0.03 ρ/kb. Hotspots had varied intensities from 2 to 20 times the background rate, set at 0.06 ρ/kb, 0.15 ρ/kb, 0.3 ρ/kb, and 0.6 ρ/kb. One scenario simulated a constant effective population size, with 500 sequences, 40 haplotypes each, with an average Watterson’s θ of 0.00355, the average between Lake Washington and Puget Sound (–nsamp 40 –len 250000 –theta 0.00355). For both populations, a bottleneck was simulated 8,000 generations ago (Puget Sound: t = 0.029, theta = 0.0036; Lake Washington: t = 0.022, theta = 0.0035). Two bottleneck strengths were simulated by setting the probability that a lineage coalesces to 10% or 90% (s = 0.1, 0.9). Overall, hotspot sharing between simulated Lake Washington and simulated Puget Sound populations was quantified by examining all pairwise comparisons between populations and bottleneck strengths. The first hotspot simulated should not be called using our method as it falls below our cutoff, but can provide information about how hotspots that fall below our cutoff affect hotspot calling. The number of false positive and false negative hotspots were calculated using custom Python scripts.

### Location of hotspots around transcription start sites

Transcript annotations from Ensembl (build 90) were lifted to the revised threespine stickleback genome assembly (Glazer et al. 2015) by aligning each transcript using BLAT (v36, default parameters) (Kent 2002). Aligned transcripts were only retained if the entire transcript aligned to the revised genome assembly. Transcript start sites (TSSs) consisted of a 2 kb region, centered at the start position of the transcript. A hotspot was considered overlapping with a TSS if the midpoint of the hotspot overlapped with any part of a 2 kb TSS region. Enrichment of hotspots in TSSs were compared against 10,000 random permutations. 2 kb regions were randomly drawn across the genome, totaling the number of hotspots identified in each population. TSS annotation filtering, overlap of hotspots with TSSs, and random permutations were completed using custom Python scripts.

### GC-Biased Substitutions

GC to AT and AT to GC substitutions were quantified within 2 kb regions of the genome that had recombination rates in the top and bottom 5% as well as within all 2 kb recombination hotspots. The top 5% of recombination rates captures regions of the genome that may broadly have high recombination rates and not contain recombination hotspots. The top 5% of recombination rates includes 96 hotspots for Lake Washington (out of 1,627 hotspots) and 314 hotspots for Puget Sound (out of 2,338 hotspots). The equilibrium GC content was calculated as the proportion of AT to GC substitutions out of the total pool of substitutions (AT to GC and GC to AT) (Meunier and Duret 2004; Singhal et al. 2015; Sueoka 1962). To increase the total number of sites available for the analysis, the ancestral allele state was inferred using only *G. wheatlandi*, rather than requiring a matching ancestral allele in both *G. wheatlandi* and *P. pungitius*. Because CpG sites can have higher mutation rates (Fryxell and Moon 2005; Weber et al. 2014), all consecutive CG sites in the ancestral sequence were removed from the analysis.

### DNA motif identification

MEME (v4.11.0) was used to identify novel DNA motifs enriched in hotspots and matched coldspots (Bailey and Eklan 1994). Each hotspot was matched to a randomly selected 2 kb coldspot, which was located at least 25 kb from any identified hotspot, contained a GC nucleotide content that was within 2% of the hotspot after removing ancestral CpG sites (GC-matched), and had a mean recombination rate that was less than half the background recombination rate of the population (Lake Washington: less than 0.017 ρ/bp; Puget Sound: less than 0.035 ρ/bp). MEME ignored motif occurrences if they were present in a hotspot multiple times (-mod zoops). This was to prevent the reporting of repetitive motifs. MEME was run separately for each chromosome and population and was completed when 50 motifs were identified (-nmotifs 50). Motif identification was conducted separately for shared hotspots and population-specific hotspots.

The DNA-binding protein, PRDM9, is important for localizing recombination hotspots in mammals (Baker et al. 2015; Baudat et al. 2010; Billings et al. 2013; Brick et al. 2012; Myers et al. 2010; Myers et al. 2008; Parvanov et al. 2010; Powers et al. 2016; Pratto et al. 2014). To determine if any PRDM genes had a role in localizing hotspots in threespine stickleback fish, FIMO (v4.11.0, default parameters) (Grant et al. 2011) was used to scan hotspot sequences for the predicted DNA binding motifs for each of the 11 annotated PRDM genes in the threespine stickleback genome (Ensembl, build 90). DNA binding motifs for each PRDM protein were predicted using the Cys**2**His**2** zinc finger prediction tool, Predicting DNA-binding Specificities for the Cys**2**His**2** Zinc Finger Proteins (Persikov et al. 2009; Persikov and Singh 2014). Predicted zinc finger domains were included if the HMMER bit score for the zinc fingers was 17.7 or higher (Persikov et al. 2009; Persikov and Singh 2014). To determine the expected number of occurrences of a motif of the same length and GC composition in hotspots, the PRDM motifs were shuffled 100 separate times. FIMO was run on the shuffled motifs to create a null distribution. Motifs were shuffled using a custom python script.

## Results

### Genetic differentiation between Lake Washington and Puget Sound

Freshwater populations of threespine stickleback fish frequently exhibit signs of past bottlenecks, consistent with their colonization from marine ancestors ~10–15 thousand years ago (Bell and Foster 1994; Ferchaud and Hansen 2016; Hohenlohe et al. 2010; Liu et al. 2016). Given the recent divergence and the close geographic proximity between Lake Washington (freshwater) and Puget Sound (marine), we first examined whether these two populations were genetically distinct. Using FastStructure, a two population model was the most highly supported (marginal likelihood = -0.834, Supplemental Figure 1).

Within each population, we explored whether there were signatures of past bottleneck events. The average nucleotide diversity within both populations was similar (Lake Washington: 0.003; Puget Sound: 0.003), whereas the genome-wide average nucleotide diversity between populations was 0.004. The nucleotide diversity values we calculated are similar to previously reported values for other marine and freshwater stickleback populations (Guo et al. 2015; Hohenlohe et al. 2010; Kitano et al. 2007). Both populations had negative Tajima’s D values (Tajima 1989), consistent with an excess of rare variants from a recent population expansion (Lake Washington: -0.422; Puget Sound: -0.723).

The demographic histories of Lake Washington and Puget Sound were estimated using Pairwise Sequentially Markovian Coalescent (PSMC) models (Figure 1). Puget Sound experienced a bottleneck from around 18,000 years ago until about 8,000 years ago where the effective population size decreased to 74,250 ± 1,259 individuals (starting N**e** = 132,700 ± 796) while Lake Washington experienced a small bottleneck around the same time where the effective population size decreased to 91,760 ± 1,960 individuals (starting N**e** = 129,138 ± 897) (Figure 1). Both populations have had a constant effective population size for the last ~5,000 years. Puget Sound has a larger effective population size than Lake Washington, matching the expected pattern of marine populations having larger effective population sizes than freshwater populations (DeFaveri and Merila 2015; Gow et al. 2006; Makinen et al. 2006).

**Figure 1.**
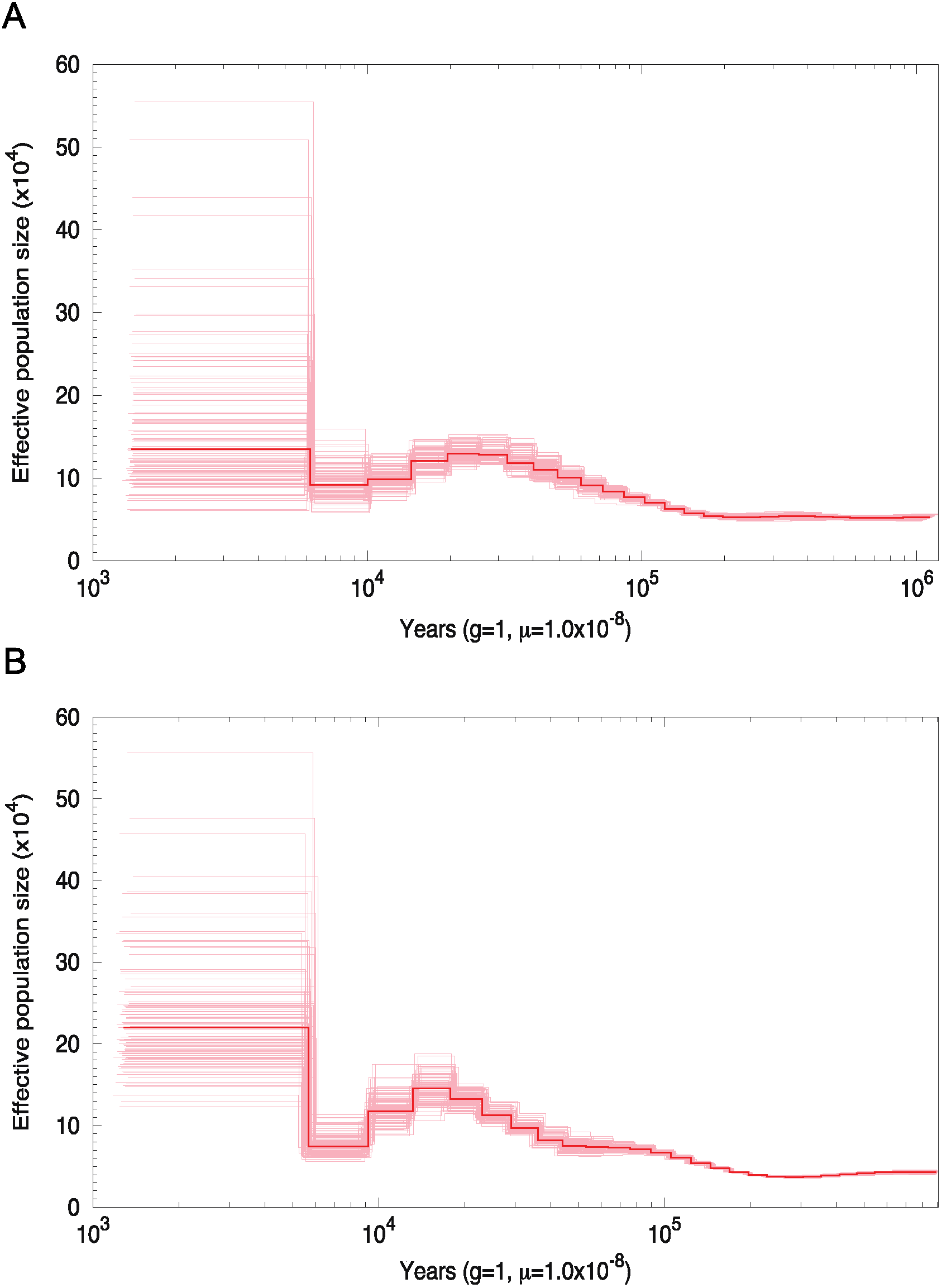
Lake Washington and Puget Sound have experienced past population bottlenecks. Demographic history for Lake Washington (A) and Puget Sound (B) was estimated using pairwise sequential Markov coalescent (PSMC) from a single female fish from each population. 100 bootstrap replicates around the estimated history are shown.

### Fine-scale estimation of recombination rates across the genome

Using a dense set of SNP markers from whole-genome sequencing, we estimated recombination rates across the genomes of Lake Washington and Puget Sound threespine stickleback fish. The average genome-wide population recombination rate in Lake Washington was half of the rate observed in Puget Sound (Lake Washington: 0.035 ρ/bp; Puget Sound: 0.072 ρ/bp; Wilcoxon Rank Test; p < 0.001, Supplemental Table 1). Despite having an overall lower genome-wide recombination rate in Lake Washington, recombination rates were largely conserved at broad scales between the two populations. We observed a highly significant positive correlation of recombination rates between the populations at the scale of 500 kb windows (Spearman’s Rank Correlation; r = 0.931, p < 0.001; Figure 2; Supplemental Figure 2). Additionally, recombination rates were lower at the center of chromosomes (center 25% of all chromosomes) and significantly higher at the ends of the chromosomes (terminal 25% of all chromosomes) for both populations (Wilcoxon Rank Test; Lake Washington: ends of chromosomes = 0.069 ρ/bp, center of chromosomes = 0.009 ρ/bp, p < 0.001; Puget Sound: ends of chromosomes = 0.108 ρ/bp, center of chromosomes = 0.016 ρ/bp, p < 0.001; Figure 2). Rate differences at chromosome ends have been documented in other populations of threespine stickleback (Glazer et al. 2015; Roesti et al. 2013; Sardell et al. 2018) as well as across a wide-range of other animals, plants, and fungi (Barton et al. 2008; Berner and Roesti 2017; Broman et al. 1998; See et al. 2006).

**Figure 2.**
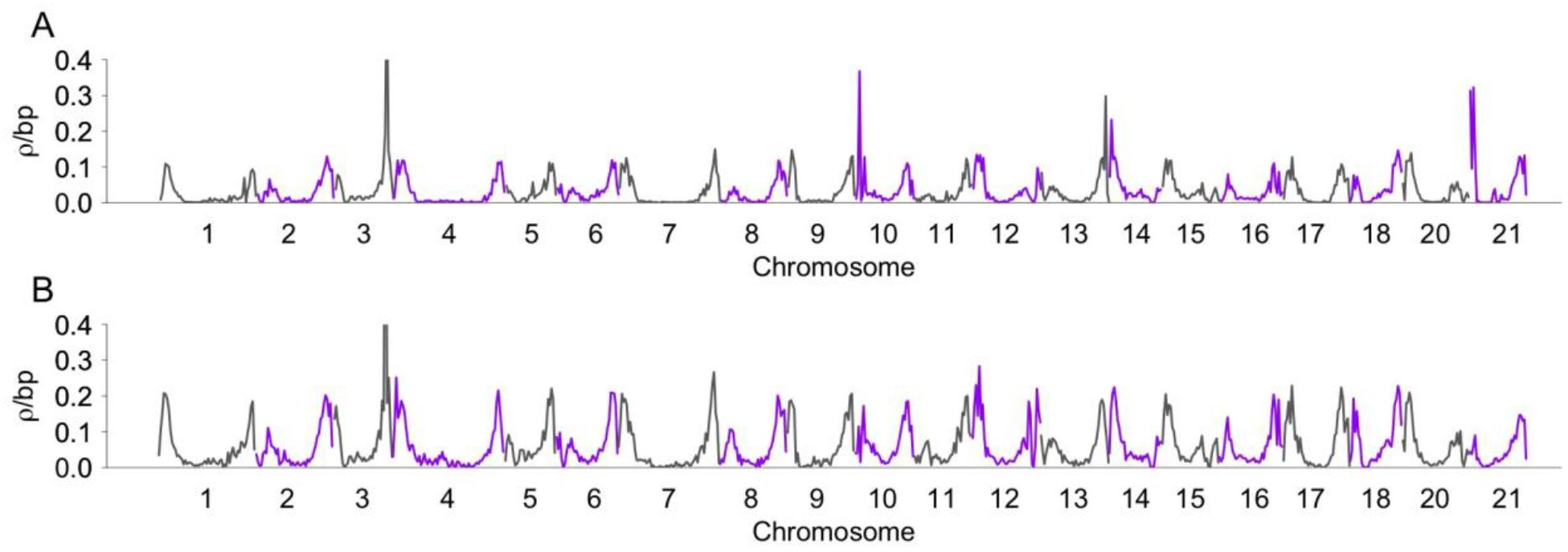
Recombination rates are similar at a broad scale in each population. Mean recombination rates were estimated using LDHelmet in non-overlapping 500 kb windows for each autosome in Lake Washington (A) and Puget Sound (B). Centromere positions are shown in Supplemental Figures 3 and 4. Transitions between gray and purple indicate different chromosomes.

To determine whether the broad-scale recombination rates we estimated from LD-based methods are concordant with recombination rates measured from linkage mapping, we compared the rates from Lake Washington and Puget Sound with the rates estimated from a genetic linkage map from a freshwater female and a marine male (Glazer et al. 2015). We found a significant positive correlation between recombination rates in both populations and the linkage map (Spearman’s Rank Correlation; Lake Washington: r = 0.830, p < 0.001; Puget Sound: r = 0.810, p < 0.001; Figure 3). These data indicate that broad-scale changes are conserved across multiple populations of threespine stickleback fish and confirm that the recombination rates estimated from LD-based methods largely parallel the rates observed from genetic linkage maps. Although broad-scale (Mb) recombination rates tend to be conserved over longer evolutionary timescales (Fledel-Alon et al. 2009; Kong et al. 2002; Serre et al. 2005; Stevison et al. 2015), fine-scale (kb) rates within chromosomes can rapidly evolve (Barton et al. 2008; Hellsten et al. 2013; McVean et al. 2004; Myers et al. 2005). In many organisms, recombination is organized locally into narrow regions of very high rates (i.e. “hotspots”), surrounded by regions of little to no recombination (i.e. “coldspots”) (Baudat et al. 2010; Jeffreys et al. 1998; Steiner et al. 2002). Consistent with this, we found highly variable fine-scale recombination rates across individual chromosomes in both Lake Washington and Puget Sound (Figure 4; Supplemental Figures 3 and 4).

**Figure 3.**
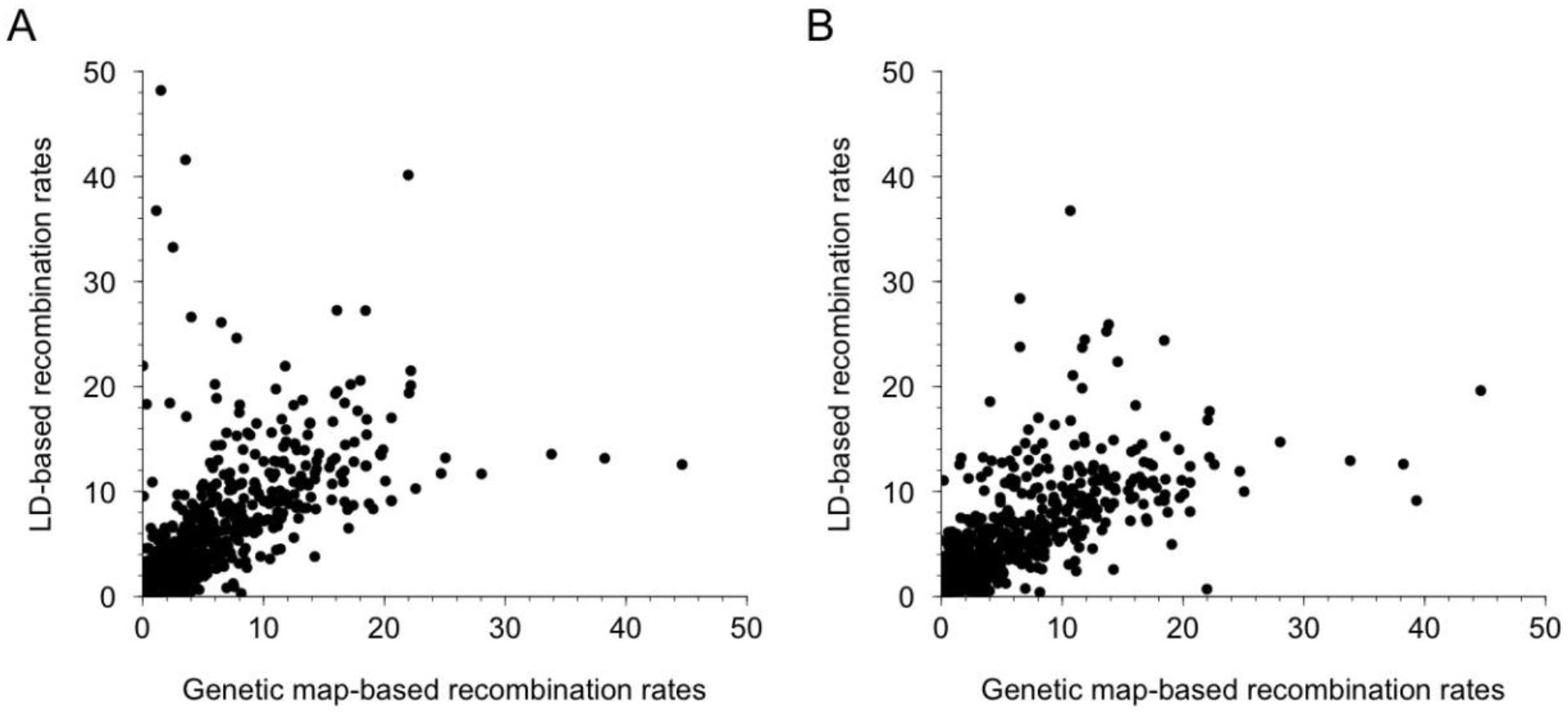
LD-based estimates of recombination rates are highly correlated with estimates from genetic linkage maps. Population-scaled recombination rates were converted to cM/Mb. There is a significant positive correlation in Lake Washington (Spearman’s rank correlation; r = 0.830; p < 0.001) (A) and Puget Sound (Spearman’s rank correlation; r = 0.810; p < 0.001) (B) between LD-based recombination rates and genetic map-based recombination rates.

**Figure 4.**
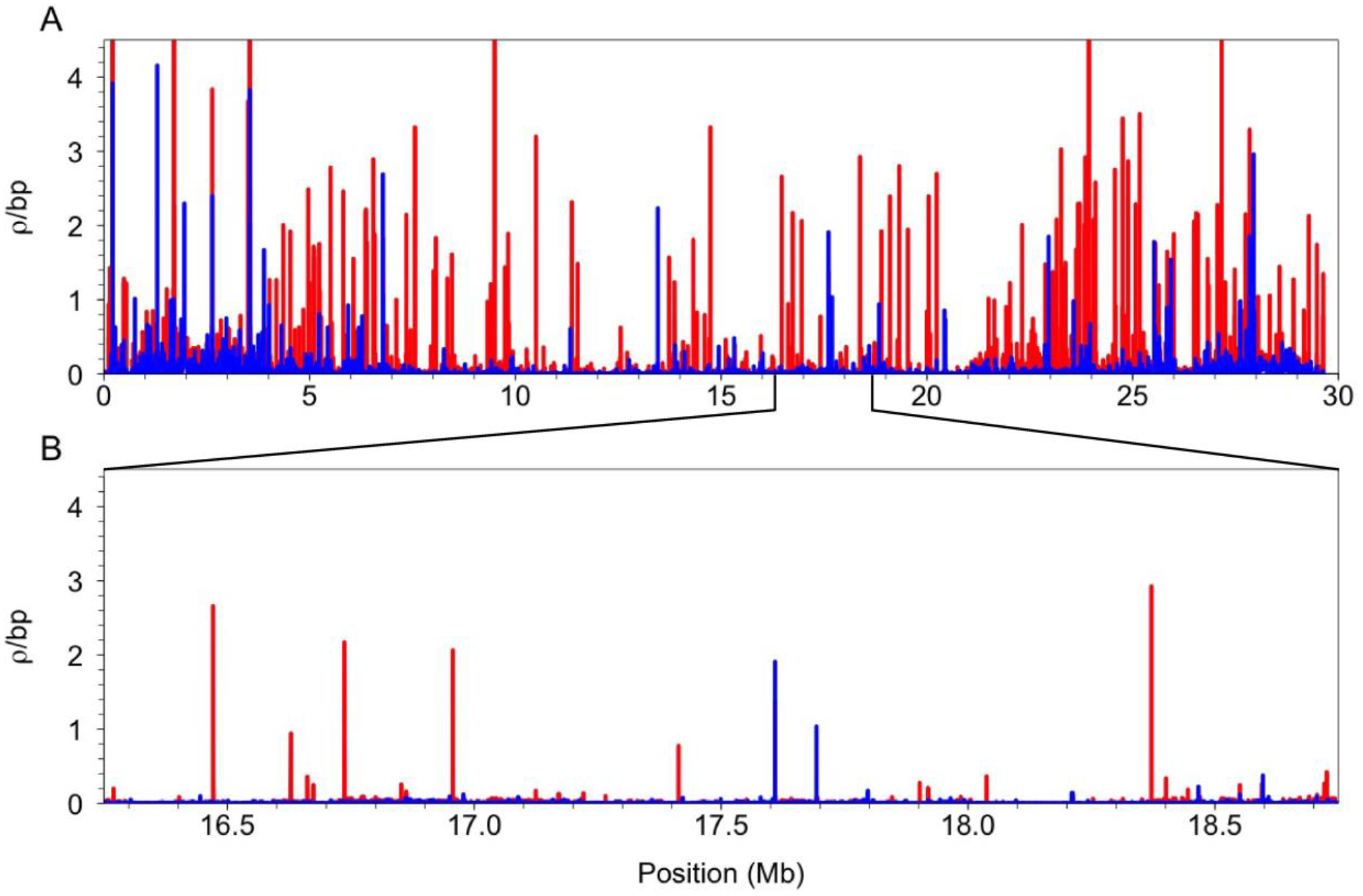
Recombination rates vary at a fine-scale across chromosome one. Population-scaled recombination rates across chromosome one are shown for Puget Sound (red) and Lake Washington (blue) (A). A subset of chromosome one is shown to highlight population-specific peaks of recombination across a narrow 2.5 Mb region (B). Only recombination rates below 4.5 ρ/bp are shown. The remaining chromosome plots are in supplemental figures 3 and 4.

### Divergent hotspot locations between populations of threespine sticklebacks

Using a sliding-window approach, we identified 2,338 hotspots in Puget Sound and 1,627 hotspots in Lake Washington. Strikingly, only 312 of these hotspots were shared between populations (13.3% of hotspots in Puget Sound and 19.2% of hotspots in Lake Washington). This lack of hotspot overlap between Lake Washington and Puget Sound may, in part, be due to hotspots falling just below the hotspot threshold. To investigate this, we looked for any increase in recombination rate in locations where hotspots were present in one population, but absent in the other. We found little evidence of a localized increase in recombination rate in these regions. Recombination rates were close to the background rate in the population where hotspots were deemed absent (Figures 5A and 5B). This pattern was even more apparent when shared hotspots were removed from the analysis (Figures 5C and 5D). The small degree of overlap we observed in hotspots between the populations was much greater than what would be expected from chance alone (10,000 random permutations; p < 0.001; Supplemental Figure 5), indicating much of the hotspot overlap likely represents shared ancestry.

**Figure 5.**
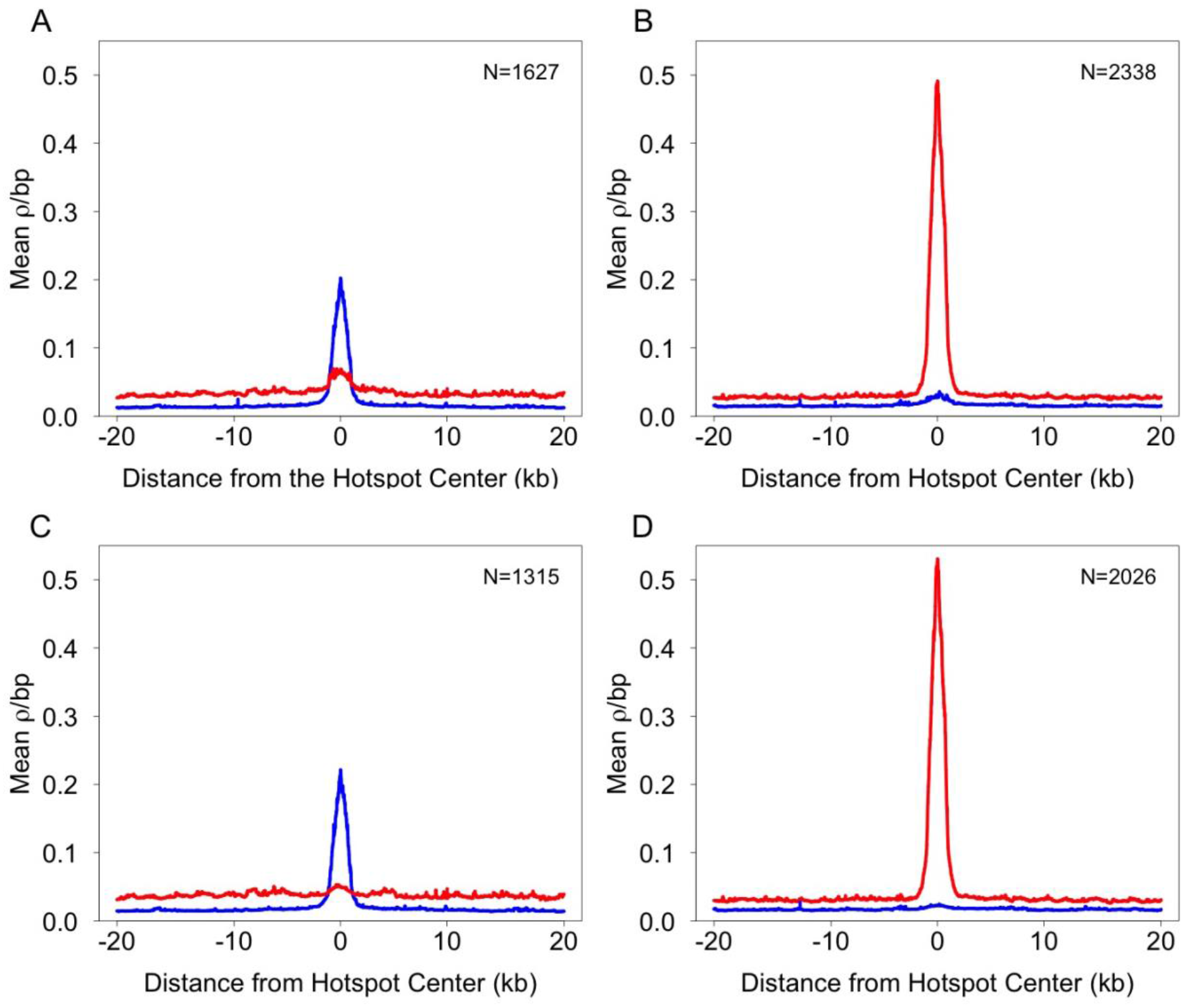
LD-based recombination rates around hotspots are population-specific. Mean recombination rates are shown across a 40 kb interval, flanking the center of hotspots. The mean recombination rate in shared and population-specific Lake Washington hotspots is higher in the Lake Washington population compared to the homologous regions in the Puget Sound population (A). The mean recombination rate in shared and population-specific Puget Sound hotspots are higher in the Puget Sound population compared to the homologous regions in the Lake Washington population (B). The pattern is more pronounced when shared hotspots are removed from the comparison, leaving only the population-specific hotspots (C and D). Puget Sound is shown in red and Lake Washington is shown in blue.

### Increased recombination rate in the pseudoautosomal region

Genetic recombination between sex chromosomes is restricted to the pseudoautosomal region (PAR), where rates of recombination can be orders of magnitude above genome-wide averages (Otto et al. 2011; Wright et al. 2016). In threespine stickleback, crossing over between the X and Y chromosomes is restricted to a ~2.5 Mb PAR (Peichel et al. 2004; Roesti et al. 2013; White et al. 2015; Yoshida et al. 2014). Because of the potential for high rates of crossing over in the PAR, we estimated population-scaled recombination rates for this region independently from the autosomes. The average recombination rate in the PAR was 0.232 ρ/bp for Puget Sound and 0.129 ρ/bp for Lake Washington. These rates were significantly higher than the average recombination rate across the autosomes (Lake Washington autosome average rate: 0.035 ρ/bp, p <0.001; Puget Sound autosome average rate: 0.072 ρ/bp, p < 0.001; Figure 6). Although we observed some fine-scale variation in recombination rates across the PAR (Figure 6), we identified very few hotspots, which may be due to the increased background recombination rate across the PAR.

**Figure 6.**
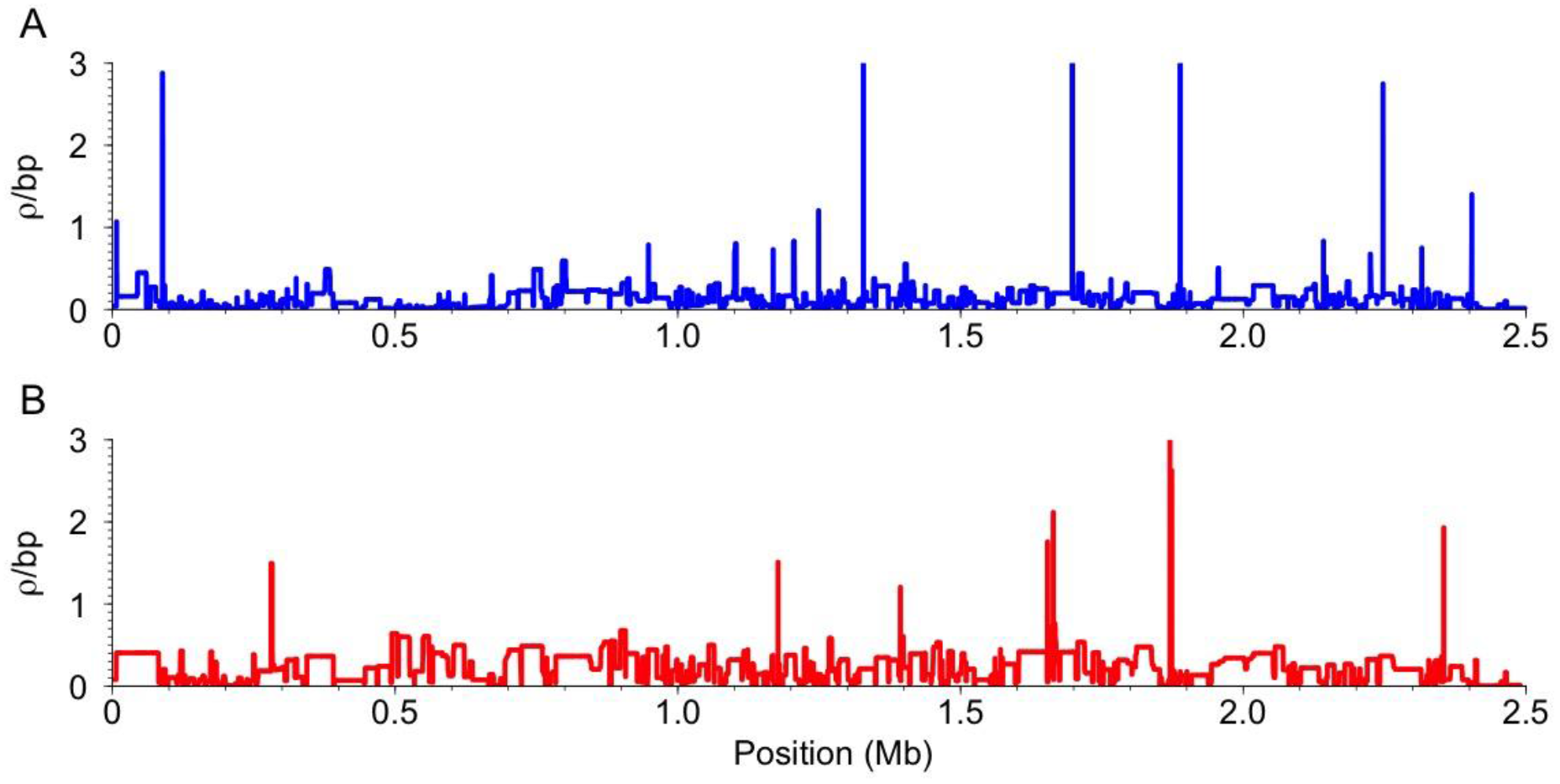
LD-based recombination rates are higher across the pseudoautosomal region (PAR). The PAR is the first ~2.5 Mb of linkage group 19. The Lake Washington (A) and Puget Sound (B) population-specific rates are shown separately. Overall, recombination rates are higher across the PAR than the autosomes (see Figure 4B) (Lake Washington PAR average: 0.129 ρ/bp; Lake Washington autosome-wide average: 0.035 ρ/bp; Puget Sound PAR average: 0.232 ρ/bp; Puget Sound autosome-wide average: 0.072 ρ/bp).

### Demographic history may not completely account for hotspot divergence

To explore how changes in past effective population size (N**e**) may have affected our ability to detect hotspots, we simulated haplotypes with known demographic histories that followed the demographic histories we estimated from Lake Washington and Puget Sound, along with a known distribution of recombination hotspots. If the minimal hotspot overlap we observed between populations of threespine stickleback fish was because of high false positive and false negative rates induced by demographic history, we would expect hotspots to be incorrectly called to a similar degree in the bottleneck simulations. Both bottleneck strengths exhibited elevated false positive and false negative rates compared to the control simulation, with the highest false positive and false negative rates under the strong bottleneck scenario (Supplemental Table 2). To determine the overall effect of elevated error rates on determining the number of shared hotspots between populations, we compared the simulated Lake Washington haplotypes to the simulated Puget Sound haplotypes from both bottleneck scenarios. Despite the elevated error rates, hotspot sharing was higher between the simulated populations than the observed number of hotspots shared between actual Lake Washington and Puget Sound populations for the weak bottleneck (weak bottleneck: Lake Washington: 59.7%; Puget Sound: 55.2%; actual Puget Sound shared hotspots: 13.3%; actual Lake Washington shared hotspots: 19.2%). This indicates that a weak bottleneck in both populations is not sufficient to drive the high degree of hotspot divergence we observed. However, if the bottleneck strength was very high (s=0.9) in both populations, elevated error rates in hotspot calling could result in a lack of hotspot overlap that mirrors the divergence we observed between populations. In this simulation, there was a similar percent of shared hotspots as observed in the actual populations (strong bottleneck: Lake Washington: 20.7%; Puget Sound: 19.8%; actual Puget Sound shared hotspots: 13.3%; actual Lake Washington shared hotspots: 19.2%).

Based on the demographic histories we estimated, Lake Washington experienced a less intense bottleneck than Puget Sound. We therefore also used simulations to explore the expected hotspot overlap if only one of the populations experienced a strong bottleneck. If Puget Sound experienced a strong bottleneck and Lake Washington experienced a weak bottleneck, 36.7% of hotspots were shared in the simulated Lake Washington population and 20.5% of hotspots were shared in the simulated Puget Sound population (actual Lake Washington shared hotspots: 19.2%, actual Puget Sound shared hotspots: 13.3%). Except for a scenario where both populations underwent a severe bottleneck in the past, our simulations suggest that demographic history alone is not sufficient to completely explain the divergence we observed in hotspot location between populations.

### Hotspots are enriched around transcription start sites

Hotspot localization in genomes varies among taxa. In yeast, birds, and some plants, where hotspots are evolutionarily conserved, hotspots tend to be enriched within transcription start sites (Kawakami et al. 2017; Pan et al. 2011; Singhal et al. 2015; Tischfield and Keeney 2012). In mammals with rapidly evolving hotspots, hotspots are typically located away from genic regions (Brick et al. 2012; Brunschwig et al. 2012; Myers et al. 2005). We investigated whether threespine stickleback fish hotspots mimic either of the patterns seen in other systems. We found an enrichment of hotspots around TSSs, compared to random permutations of hotspots (Lake Washington: 26% of hotspots fell within 3 kb of a TSS, p < 0.034; Puget Sound: 29% of hotspots fell within 3 kb of a TSS, p < 0.001; Supplemental Figure 6). This pattern also held when examining only population specific hotspots (Lake Washington: p = 0.007; Puget Sound: p < 0.001; Supplemental Figure 7); however, shared hotspots were not enriched in TSSs compared to random permutations (Lake Washington: p = 0.370; Puget Sound: p = 0.827; Supplemental Figure 6). The lack of significant enrichment of shared hotspots around TSSs is likely due to the small sample size. When we randomly drew samples from the population-specific hotspots that were equal in size to the shared hotspot pools, there was no longer enrichment around TSSs (Lake Washington: p = 0.947; Puget Sound: p = 0.808).

### Regions of high recombination exhibit GC-biased nucleotide substitution

Recombination leaves distinct signatures of nucleotide substitution across the genome (Duret and Arndt 2008; Mugal et al. 2015; Webster and Hurst 2012). Over time, the repair of heteroduplex DNA during meiosis favors the substitution of GC nucleotides over AT nucleotides, which increases the frequency of GC nucleotides, leading to GC-biased base composition (Lesecque et al. 2013; Marais 2003; Meunier and Duret 2004). Regions of the genome with higher recombination rates tend to have higher GC-biased base composition (Kawakami et al. 2017; Kong et al. 2002; Meunier and Duret 2004; Singhal et al. 2015). To determine whether regions of higher recombination rate showed signatures of GC-biased gene conversion, we calculated equilibrium GC content (Meunier and Duret 2004; Singhal et al. 2015; Sueoka 1962) in regions of the genome with the highest and lowest recombination rates (top and bottom 5%) as well as within recombination hotspots.

In both Lake Washington and Puget Sound, we detected a significantly higher equilibrium GC content in regions of the genome with a high recombination rate (top 5% of recombination rates among 2 kb windows) compared to regions of the genome with recombination rates in the bottom 5% (Table 1). Overall, these results indicate GC nucleotide composition is influenced by the historical recombination landscape across the threespine stickleback genome. Interestingly, although hotspots in both populations have locally elevated recombination rates, there was not a parallel increase in equilibrium GC content. Equilibrium GC content in population-specific hotspots and shared hotspots was not significantly different than regions of the genome with the lowest recombination rates (bottom 5%) (Table 1). Our results are consistent with recombination hotspots being more recently derived, where locally increased recombination rates have not yet had an effect on GC-biased nucleotide substitution.

**Table 1.**
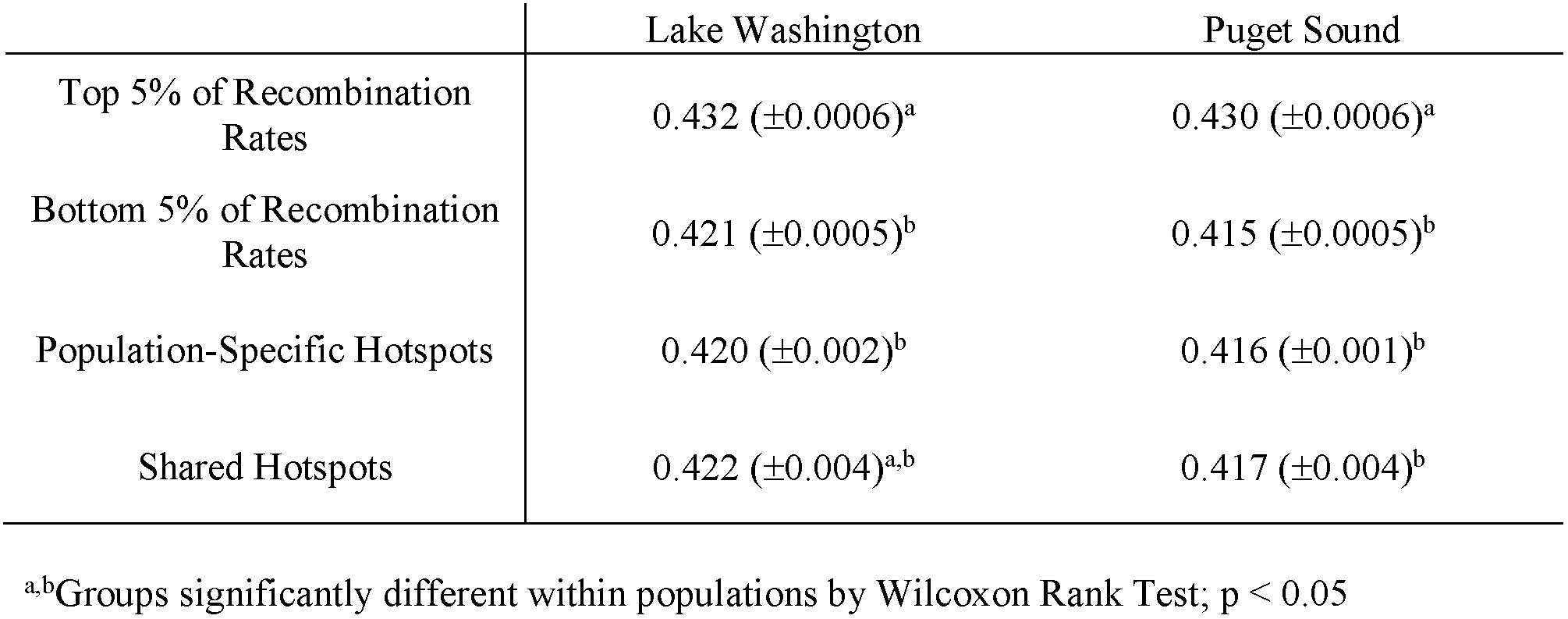
Mean equilibrium GC content (± SE)

### PRDM genes are weakly associated with threespine stickleback recombination hotspots

Hotspots in many species are targeted to specific regions of the genome by DNA binding motifs (Baudat et al. 2010; Kon et al. 1997; Myers et al. 2008; Steiner et al. 2002). In species where PRDM9 targets recombination hotspots to specific regions of the genome, the zinc finger domain of PRDM9 is typically under strong positive selection (Baker et al. 2015; Baudat et al. 2010; Billings et al. 2013; Myers et al. 2010; Oliver et al. 2009; Pratto et al. 2014) and the protein contains functional KRAB and SSXRD domains (Baker et al. 2017). In Teleost fish, two paralogs of PRDM9 have been identified, PRDM9α which contains all the protein domains and PRDM9β which lacks the KRAB and SSXRD domains (Baker et al. 2017). Threespine stickleback fish appear to have lost PRDM9α, but retain PRDM9β without the SSXRD and KRAB domains. Consistent with a lack of function directing recombination hotspots, we did not observe strong signatures of positive selection in the zinc finger domain of PRDM9β. We found zero fixed differences between threespine and blackspotted stickleback for the PRDM9 ortholog. There was one synonymous and one nonsynonymous mutation at moderate frequency in Lake Washington and two synonymous and three nonsynonymous mutations at moderate frequency in Puget Sound, indicating these mutations are likely not causing the population-specific localization of hotspots we observed between Lake Washington and Puget Sound.

We also examined whether the predicted binding sites of any of the 11 previously annotated PRDM genes in threespine stickleback fish were enriched in recombination hotspots. Less than 14% of hotspots contained any of the predicted PRDM zinc finger binding domain motifs (Supplemental Table 3). However, six of the motifs were significantly enriched in hotspots, including PRDM9, when compared to scrambled motifs of the same size and GC content (Supplemental Table 3), indicating PRDM genes could have some role in localizing a subset of recombination hotspots. Outside of PRDM9 in mammals, multiple DNA binding motifs assist with hotspot targeting in other systems such as *Schizosaccharomyces pombe* (Kon et al. 1997; Steiner et al. 2002). To see if other DNA motifs were targeting hotspots in threespine stickleback fish, we searched for motifs enriched in hotspots. The most significant motifs identified were simple mono- or di-nucleotide repeats which were present only in a subset of the hotspots (Supplemental Figure 8). These repeats were not specific to hotspots as they were also found in GC-matched coldspots.

## Discussion

### Broad-scale recombination rates across the threespine stickleback genome

At a broad scale, recombination rates across the threespine stickleback genome were conserved between the two populations. This broad scale conservation of recombination rates is a feature observed in many taxa (Fledel-Alon et al. 2009; Kong et al. 2002; Serre et al. 2005; Stevison et al. 2015) and may reflect the necessity of crossing over for the proper segregation of chromosomes during meiosis (Davis and Smith 2001; Fledel-Alon et al. 2009; Kaback et al. 1992; Mather 1936). Additionally, we observed differential rates of recombination associated with broad genomic regions that have been observed in other systems. First, we observed higher recombination rates towards the telomeres. In many species, the ends of chromosomes have higher rates of recombination (Barton et al. 2008; Berner and Roesti 2017; Kong et al. 2002; Roesti et al. 2013; Sardell et al. 2018), which is thought to be driven by male-specific localization of recombination (Broman et al. 1998; Moen et al. 2008; Singer et al. 2001). Our LD-based method estimates sex-averaged recombination rates, which does not allow us to test whether the pattern we observed around the ends of chromosomes is driven by males. However, sex-specific genetic linkage maps between the Japan Sea stickleback *(Gasterosteus nipponicus)* and the threespine stickleback *(G. aculeatus)* corroborate this pattern (Sardell et al. 2018). Second, we observed higher recombination rates in the pseudoautosomal region compared to the autosomes. Recombination rates in pseudoautosomal regions are often orders of magnitude above autosome-wide averages, as an obligate crossover should occur between the X and Y chromosomes in these small regions during every male meiosis (Hinch et al. 2014; Kauppi et al. 2012; Otto et al. 2011).

Overall, the genome-wide average recombination rate for Puget Sound was two-fold higher than in Lake Washington. Rate variation between populations or species can be driven by a number of processes. Structural variation (i.e. inversions, chromosomal rearrangements, and copy number variants) can contribute to rate variation among genomes. Indeed, recombination rates have been shown to vary across chromosomal regions due to segregating inversions between marine and freshwater populations of threespine stickleback (Glazer et al. 2015; Jones et al. 2012). Although structural variants could explain rate differences between Lake Washington and Puget Sound populations at a more localized level, they cannot explain the genome-wide rate differences we observed. Over longer evolutionary timescales, recombination rate also can evolve neutrally (Dumont and Payseur 2008), driving genome-wide rate variation between species. However, neutral divergence is likely not occurring at a pace that would alter genome-wide recombination rates between recently diverged populations of threespine stickleback fish. One plausible explanation for the observed rate differences is differences in demographic history between the Lake Washington and Puget Sound populations. A larger effective population size could increase the population-scaled recombination rate (Burt 2000; Charlesworth 2009). In threespine stickleback, marine populations typically have a larger N**e** than freshwater populations (DeFaveri and Merila 2015; Gow et al. 2006; Makinen et al. 2006), consistent with our observed pattern of a higher recombination rate in Puget Sound relative to Lake Washington.

### Identifying hotspots using patterns of linkage disequilibrium

LD-based estimates of recombination rates can be affected by demographic processes that change patterns of linkage disequilibrium across the genome (Chan et al. 2012; Dapper and Payseur 2017; Johnston and Cutler 2012; McVean et al. 2004; Wall and Stevison 2016). The duration and timing of these events can have varying effects on hotspot identification, often reducing the power to detect hotspots and increasing the rate of errors (Dapper and Payseur 2017). Threespine stickleback fish have a complex history of bottleneck events and population expansions over the last 10–15 thousand years which vary across geographic regions (Bell and Foster 1994; Ferchaud and Hansen 2016; Hohenlohe et al. 2010; Liu et al. 2016; Orti et al. 1994). Based on simulations, demographic history likely has some role in the observed divergence in hotspot location between Lake Washington and Puget Sound populations, but it seems likely that population demography does not completely explain the pattern. Only in the scenario where both populations experienced a strong bottleneck do error rates rise high enough to mimic the observed divergence in hotspot location. However, our estimates of effective population size over time revealed that Lake Washington and Puget Sound did not experience similar fluctuations. Both populations began with effective population sizes that largely parallel those observed in other threespine stickleback fish populations (Liu and Hansen 2017; Ravinet et al. 2018). Puget Sound then experienced a larger population expansion roughly 18,000 years ago, followed with a decrease in population size at approximately 8,000 years ago. Lake Washington had a slight increase in population size, followed by a small bottleneck around the same time, but overall changes in effective population size were more stable in this population. Examination of where recombination hotspots are currently forming across the genome in Lake Washington and Puget Sound would help confirm the patterns we observed. Surveys of double strand break hotspots (Pratto et al. 2014; Smagulova et al. 2011) or crossover breakpoints in genetic crosses (Broman et al. 1998; Campbell et al. 2016; Drouaud et al. 2006; Marand et al. 2017) would reveal the degree to which recombination hotspots are targeted to different genomic locations in these two populations.

### Hotspot evolution in freshwater and marine threespine stickleback populations

Of the 3,965 hotspots between Lake Washington and Puget Sound, only ~15% of hotspots are shared, indicating many of the hotspots are recently derived within populations of threespine stickleback fish. Consistent with the recent evolution of hotspots, we did not observe an elevated equilibrium GC content in these regions. One possible model is that recombination hotspots can shift over short evolutionary timescales among regions of the genome that are susceptible to homologous recombination, such as regions of accessible chromatin. Both evolutionarily conserved and rapidly evolving hotspots tend to locate to regions of accessible chromatin (Lam and Keeney 2015; Ohta et al. 1994; Pan et al. 2011; Tischfield and Keeney 2012) or regions with histone 3 lysine 4 trimethylation (H3K4me3) (Auton et al. 2013; Baker et al. 2015; Marand et al. 2017; Smagulova et al. 2011).

In taxa where hotspots are evolutionarily conserved, hotspots are highly enriched around TSSs (Auton et al. 2013; Kawakami et al. 2017; Pan et al. 2011; Singhal et al. 2015; Tischfield and Keeney 2012).This pattern could be due to either higher selective constraints at TSSs or the chromatin structure at TSSs. TSSs are often under purifying selection and if a genomic feature, like a DNA motif, is targeting hotspots to these regions, these features would also be preserved through purifying selection, maintaining the location of the hotspot (Kawakami et al. 2017; Lam and Keeney 2015; Singhal et al. 2015; Tsai et al. 2010). On the other hand, an open chromatin conformation could be driving this pattern. TSSs and the surrounding regions must be accessible for transcription to occur while also providing sites for Spo11 to bind, initiating recombination (Lee et al. 2004; Pokholok et al. 2005) as Spo11 will create double strand breaks at any sites with accessible chromatin (Celerin et al. 2000; Ohta et al. 1994; Pan et al. 2011). In Lake Washington and Puget Sound populations, we found some enrichment of hotspots at TSSs (Lake Washington: 26% of hotspots fell within 3 kb of a TSS; Puget Sound: 29% of hotspots fell within 3 kb of a TSS) which is similar to hotspot enrichment around TSS in taxa that do not have a functional PRDM9 protein. In birds and dogs, for example, ~20–30% of hotspots overlap with TSSs (Auton et al. 2013; Kawakami et al. 2017; Singhal et al. 2015). Additional characterization is needed to determine if hotspots in threespine stickleback are occurring in regions of the genome that are already open due to transcription or if there is a mechanism that creates accessible chromatin specifically for double strand break formation, like what is believed to occur with PRDM9 in mammalian species (Diagouraga et al. 2018; Hayashi et al. 2005; Powers et al. 2016).

In some mammalian systems, positive selection acting on the zinc finger binding domain of PRMD9 has led to multiple distinct DNA binding motifs between closely related species (Baudat et al. 2010; Myers et al. 2010; Myers et al. 2008; Pratto et al. 2014). This leads to a rapid evolution of hotspot localization (Baker et al. 2015; Brick et al. 2012; Pratto et al. 2014; Smagulova et al. 2016; Stevison et al. 2015). Typically, ~40% of hotspots will contain a PRDM9 motif in mouse and humans (Baudat et al. 2010; Myers et al. 2008). In threespine stickleback fish, we found that less than 14% of hotspots had PRMD9 or any other PRDM motifs, contrary to what we would expect if PRDM9 was controlling hotspot location in threespine stickleback.

In addition, we did not find any other DNA motifs enriched in hotspots that would indicate a role of an alternative DNA-binding protein that could localize hotspots. Additional characterization is needed to understand what genomic features could be targeting hotspots, leading to the distinct fine-scale recombination landscapes observed between populations of threespine stickleback fish.

## Acknowledgements

This work was supported by the Office of the Vice President of Research at the University of Georgia (M.A.W); and the National Institute of Health (T32GM007103 to A.F.S). We thank Robert Schmitz for help with library preparations and Catherine Peichel for providing population samples. Sequencing was conducted at the Georgia Genomics and Bioinformatics Core at the University of Georgia. Raw sequences are deposited in NCBI’s Short Read Archive, reference number SRP137809 (https://submit.ncbi.nlm.nih.gov/subs/sra/SUB3748706/overview) and custom scripts can be found on Open Science Framework under A.F.S. (https://osf.io/dezug/).

